# Subchronic oral toxicity study of Aldicarb sulfoxide in Sprague-Dawley rats

**DOI:** 10.1101/2022.09.29.510057

**Authors:** Yongchao Ji, Yi Liu, Juanjuan Duan, Yiting Wang, Yu Wang, Fan Wang, Chao Chen, Wensheng Zhang

## Abstract

Aldicarb sulfoxide is the metabolite of the exceedingly deadly carbamate insecticide Aldicarb. Aldicarb sulfoxide is easily soluble in water and has been identified in groundwater. Moreover, Aldicarb sulfoxide is highly hazardous to mammals. However, the toxicity data of Aldicarb sulfoxide is scarce. Here, the effects of relatively low dosages of exposure to Aldicarb sulfoxide on the blood biochemical, hematological, and histopathological parameters were examined in SD rats. The rats were treated with Aldicarb sulfoxide orally at dosages of 0, 6.3, 18.9, and 56.7 μg per kg b.w. daily for 12 weeks. The results revealed that Aldicarb sulfoxide induced significant alterations in liver-related biochemical parameters, glucose level, cholesterol level, and uric acid level. Histopathological testing demonstrated cellular alterations in the liver of 18.9 and 56.7 μg/kg BW. Together, our data demonstrated that Aldicarb sulfoxide exposure causes liver injury. Furthermore, the no observed adverse effect level (NOAEL) was less than 6.3 μg/kg BW, and the lowest benchmark dose (BMD) value was 0.03 μg/kg BW. Accordingly, Aldicarb sulfoxide’s derived no-effect level (DNEL) was calculated as 0.00035 μg/kg BW. Thus, this work is an important reference for supervising Aldicarb sulfoxide in the environment and determining safe exposure limits for Aldicarb sulfoxide.

## 1. Introduction

Aldicarb sulfoxide (ASX), is the common name for 2-methyl-2-(methylsulfonyl) propionaldehyde O-(methylcarbamoyl) oxime, represented by the molecular formula C_7_H_14_N_2_O_3_S. It is the metabolite of Aldicarb [2-methyl-2-(methylthio) propionaldehyde O-(methylcarbamoyl) oxime], which is an oxime carbamate pesticide widely used for crop protection. The metabolites of Aldicarb are the active components of the pesticide. Aldicarb is applied in granule form and released by soil moisture; immediately, it starts to be oxidised into ASX, some of which is further oxidised (Lightfoot et al., 1987). Aldicarb and its residues are highly water soluble; the residues found in drinking water are generally present in a 1:1 ratio of sulfoxide to sulfone (DePass et al., 1985). The half-lives of ASX and Aldicarb sulfone (ASN) mixtures are under three years, and the reaction rate constant for the two compounds are relatively constant below pH 7 and increase rapidly as the PH increase beyond 7 (Hansen et al., 1983). In the United States, ASX was detected in groundwater in the range of 0.01-1030.00 μg/L (EPA, 1992), and a reference dose of 1 μg/kg BW was given by EPA (EPA, 1995). However, the acute reference dose (RfD) given by EPA was calculated based on the toxicity data of Aldicarb. ASX is a highly acutely toxic metabolite; rat oral LD50 is 490 μg/kg BW (PubChem, https://pubchem.ncbi.nlm.nih.gov/). Therefore, ASX may be a considerable risk to human health. ASX is highly toxic to Daphnia laevis, EC50 values are near or below 50 μg/L (Jeffery et al., 1985). Approximately 1800 μg/kg body weight/day of a 1:1 solution of ASX/ASN in drinking water exposure to rats for 29 days can cause substantial effects (DePass et al., 1985). To the best of our knowledge, few ASX toxicity data are available. As such, it is imperative to characterise the ASX toxicity and assess its health risk.

The current study was conducted in Sprague-Dawley rats after 12 weeks of oral administration of various doses of ASX. To assess the toxic response, blood and tissues (heart, liver, lung, spleen, kidney, brain, adrenal gland, testis, epididymis, uterus, ovary, thymus, lymph) were collected and analyzed for haematology, serum biochemistry, and histopathology. The lower level of the BMD’s 90% confidence interval (BMDL) was then calculated to determine the DNEL.

## 2. Materials and Methods

### 2.1. Chemicals

Aldicarb sulfoxide (≥ 98.0% ASX, Sigma-Aldrich-Chemie, Steinheim, Germany). All reagents and chemicals were of analytical grade quality or higher purity. ASX was dissolved in ddH_2_O.

### 2.2. Animals

Healthy 6-week-old SD rats were purchased from Beijing Vital River Laboratory Animal Technology Co., Ltd. (Beijing, China). A total of 80 animals (5 rats/cage) were housed under the standard controlled conditions (temperature (20-24 □), light (12 h night:12 h day)) relative humidity of 35% to 60%. During the experiment, they were allowed free access to standard rat chow (purchased from Beijing HFK Bioscience Co., Ltd (Beijing, China)) and tap water. After 5 days of acclimation, rats were weighed and randomized into experimental and control groups. All animal procedures were conducted by the Guidance for the Care and Use of Laboratory Animals (Clark et al., 1997) and were approved by the Institutional Animal Use and Care Committee of Beijing Normal University.

### 2.3. Study Design and Experiment Procedure

Rats were randomly divided into groups (10/sex/dose): one control group and three treated groups. Twenty rats were used per group, by the OECD Test guideline TG408 “Repeated Dose 90-day Oral Toxicity Study in Rodents” (OECD, 2017). The initial concentration of ASX was used in low, middle, and high exposure groups (6.3, 18.9, and 56.7 μg per kg BW). Treatment of all animals was performed by oral gavage (0.5 mL per 100 g) for 12 weeks. In the tenth week, two male rats in the high-dose group died of fighting. Animals were exposed continuously until the time of sacrifice, being removed from treatment only for bleeding.

### 2.4. Hematological and blood biochemical analysis

Rat heart whole-blood was placed in a centrifuge tube containing EDTA-Na_2_ for immediate analysis. The white blood cell (WBC) count, red blood cell (RBC) count, hemoglobin (HGB), platelet (PLT) count, platelet distribution width (PDW), mean platelet volume (MPV), platelet packed volume (PCV), large platelet ratio (P-LCR), percent neutrophils (NEUT%), neutrophils absolute value (NEUT), percent lymphocytes (LY%), lymphocytes absolute value (LY), hematocrit (HCT), percent monocytes (MONO%), monocytes absolute value (MONO), percent eosinophil (EOS%), eosinophil absolute value (EOS), percent basophil (BASO%), basophil absolute value (BASO), mean red blood cell volume (MCV), mean hemoglobin account (MCH), mean hemoglobin concentration (MCHC), red blood distribution width-coefficient of variation (RDW-CV) and red blood cell volume distribution width-standard deviation (RDW-SD) were analyzed by a BC6600PLUS automated blood analyzer (Mindray, China).

Rat heart whole-blood was collected in vacuum blood collection tubes without anticoagulant and centrifuged at 3000 rpm for 10 min to obtain serum samples. Serum alanine aminotransferase (ALT), aspartate aminotransferase (AST), alkaline phosphate (ALP), total protein (TP), albumin (ALB), globulin (GLB), ratio of white balls (A/G), urea (UREA), creatine (CREA), total cholesterol (CHOL), total bilirubin (TBIL), direct bilirubin (DBIL), indirect bilirubin (IBIL), cholinesterase (CHE), gamma-glutamyl transpeptidase (GGT), uric acid (UA), glucose (GLU), triglyceride (TG), high-density lipoprotein cholesterol (HDLC), low-density lipoprotein cholesterol (LDLC), creatine kinase (CK), lactate dehydrogenase (LDH), serum potassium (K), serum sodium (Na), serum calcium (Ca), serum magnesium (Mg), serum phosphorus (P), serum chloride (Cl) and serum iron (Fe) were detected using an AU5800 automatic biochemical analyzer (Beckman Coulter, USA).

### 2.5. Histopathological analysis

The extracted organs were embedded in paraffin after being fixed in 10% neutral-buffered formalin. Hematoxylin and eosin were used to stain the embedded tissue blocks, which were cut into 3-m thick sections. The stained sections were examined under a light microscope (Axio Scope A1, 07745 Jena, Germany), and the nomenclature of Roger et al. (2009) was used for histopathological assessments.

### 2.6. Body and organ weight

At the end of the experiment, final body weights and relative body weight gain (RBWG) for all animals were calculated and recorded. RBWG was calculated by the following equation (Baralić et al, 2020):

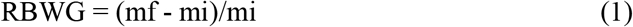

In this equation, mf signifies final body weight, while mi signifies initial body weight. The organs were collected and weighed, and the relative organ weight was calculated by dividing the organ weight by the final body weight.

### 2.7. Statistical analysis

All data were expressed as the mean ± SEM and analyzed with python software (https://www.python.org). When the variance was homogeneous, one-way ANOVA with the LSD test was used to determine the statistical significance of differences between the experimental groups and the control. When this requirement was not met, the differences between groups were compared using the nonparametric Kruskal Wallis method and the Steel Dwass test. A *p-value* of less than 0.05 was considered to be statistically significant. In addition, the benchmark dose (BMD) was calculated using online PROAST software version 70.0 (https://r4eu.efsa.europa.eu/).

## 3. Results

### 3.1. Body weight gain, food, and water consumption

Tables S1 and S2 show the RBWG and percentage difference in RBWG of rats after 12 weeks of oral ASX exposure. During the experiment, rats’ body weight was measured once a week. Male rat body weight gain in the middle dose group was 10% higher than in the control group from the first to the eleventh week, and in the high dose group from the eighth to the end of the experiment. However, an adverse effect on body weight or body weight gain is defined as any weight above or below 10% of the control value in the WHO Guidance document (Pesticide Residues in Food) of the WHO Core Assessment Group on Pesticide Residues (WHO, 2015). According to this criterion, because the majority of the observed changes in the RBWG in the middle and high dose groups are greater than 10% after the eighth week of ASX treatment, they should be considered adverse. There is no difference between the female rat and the control. There was no discernible difference in the exposed rats’ daily food and water intake (data not shown).

### 3.2. Relative organ weight

Relative organ weights (ROW) are presented in Table S3 and S4. There was no significant difference in ROW in female rats exposed to ASX compared to that in the control. While there was a significant increase in heart relative weight in the low dose group in comparison with the control (Table S4).

### 3.3. Hematological parameters

Hematological parameters measured in the present study are presented in Table S5 and S6. In female rats receiving low-dose ASX, a statistically significant increase in MCHC and RDW-CV was noted compared to the control. However, not toxicologically significant as the increases are relatively small and not dose related. In male rats receiving low-dose ASX, the WBC, NEUT, and MCHC were elevated compared to the control.

### 3.4. Serum biochemistry parameters

Serum biochemistry parameters after 12 weeks of exposure to the investigated substance are presented in Table S7 and S8. Changes in serum biochemical parameters in female and male rats are presented in Fig. 1 and 2, respectively. In the female rats, cholinesterase (CHE) activity in the three treated groups was statistically significantly decreased compared to the control (Fig. 1A), with a reduction of 9.12%, 9.94%, and 11.63%, respectively. There was a significant decrease in the total cholesterol level in the three treated groups compared to the control (Fig. 1 B). A significant decrease in the low-density lipoprotein level was noted only in the low-dose group and high-dose group. A significant elevation of the uric acid was noted only in the high dose group compared to the control (Fig. 1 C). In the male rats, CHE activity in the three treated groups was statistically significantly decreased compared to the control (Fig. 2 A), with a reduction of 23.72%, 20.31%, and 28.12%, respectively, while a significant elevation of the direct bilirubin level in the three treated groups was noted compared to the control (Fig. 2 B). There was a significant increase in glucose (GLU) only in the high dose group compared to the control (Fig. 2 C).

**Fig. 1.**
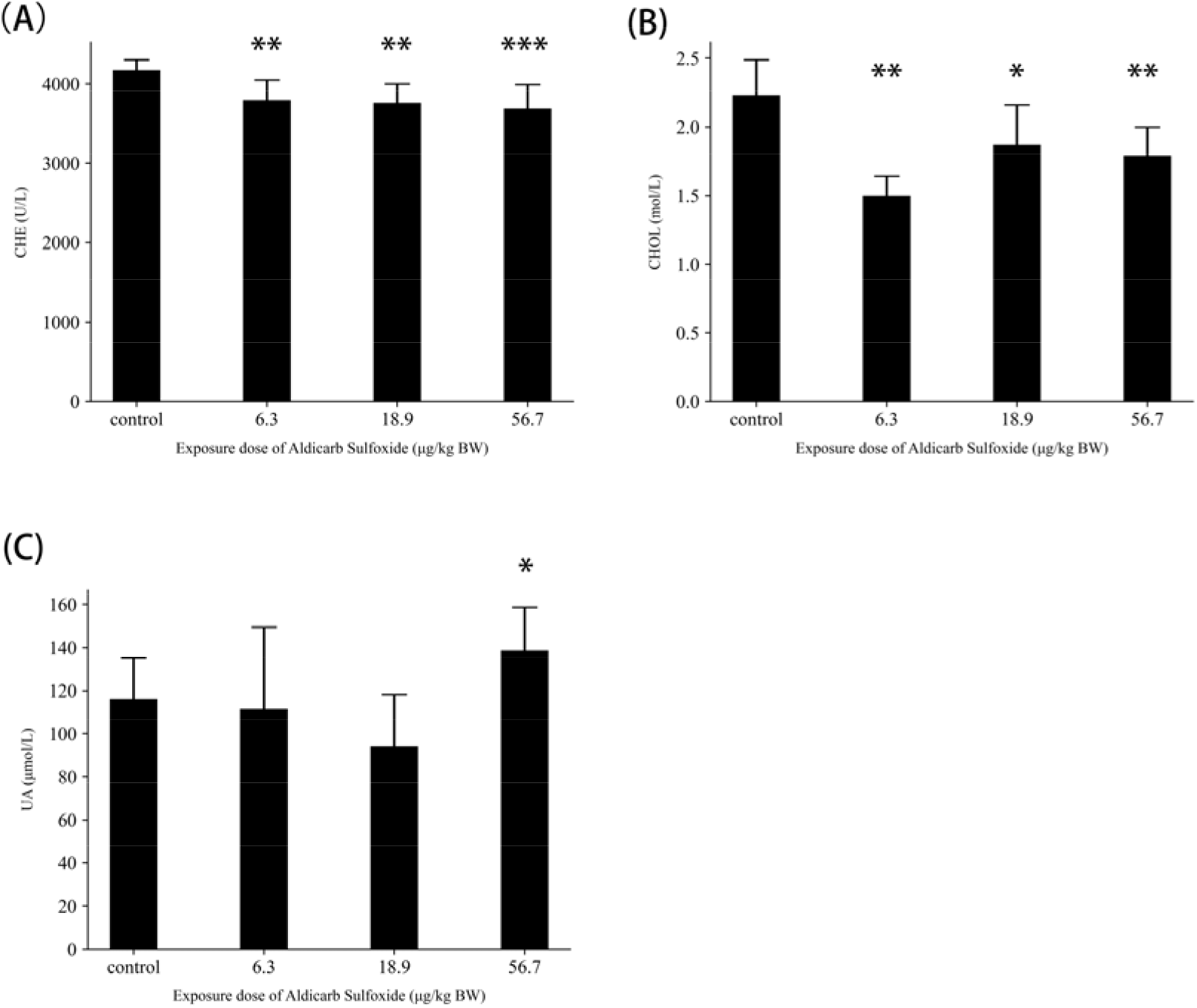
Changes in serum biochemistry parameters in the female rats after 12 weeks of oral exposure to ASX (0, 6.3, 18.9, and 56.7 μg/kg BW). (A) CHE activity (U/L), (B) the total cholesterol concentration (mol/L), (C) the uric acid concentration (μmol/L) (* p < 0.05, ** p < 0.01(compared to the control)).

**Fig. 2.**
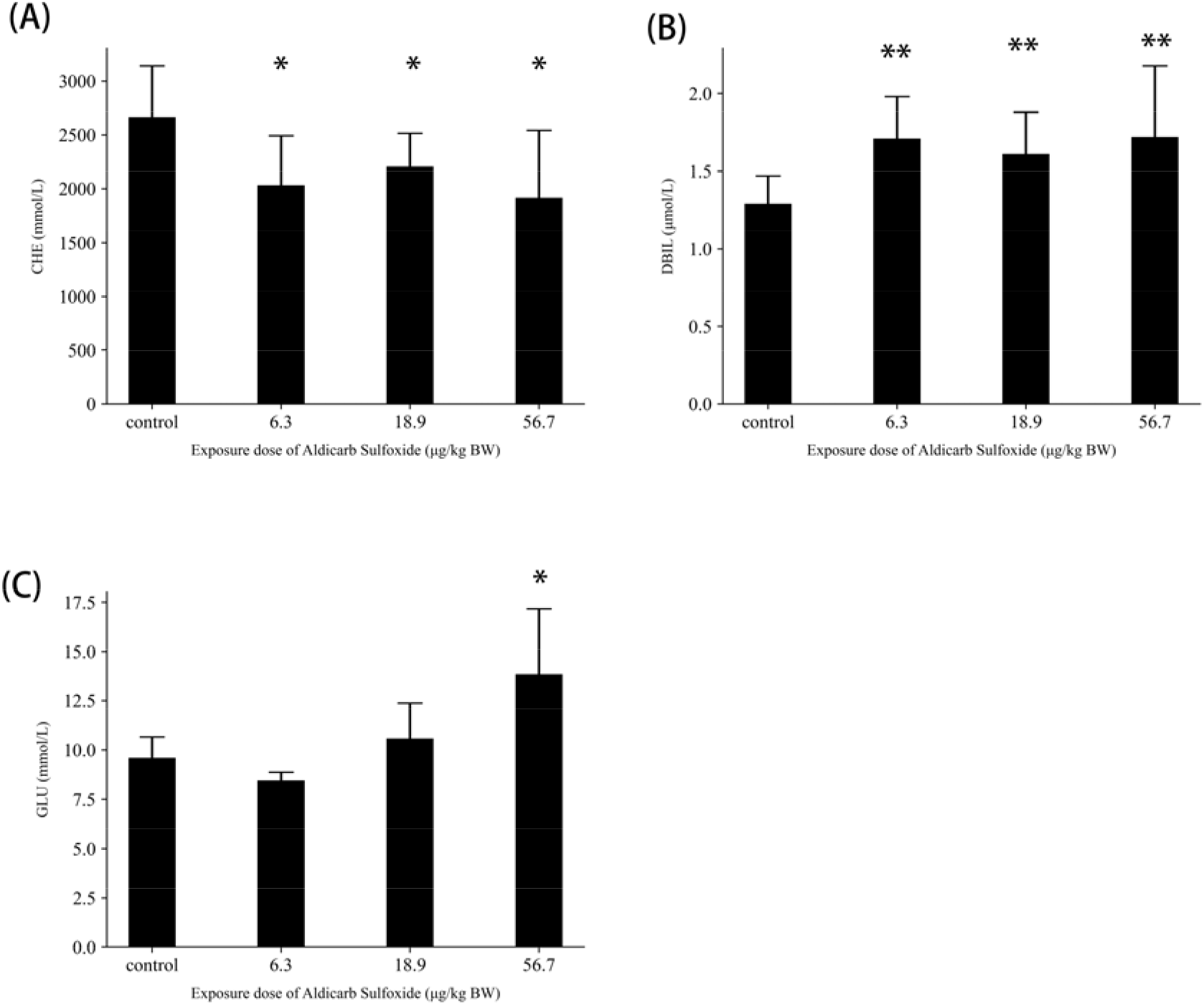
Changes in serum biochemistry parameters in the male rats after 12 weeks of oral exposure to ASX (0, 6.3, 18.9, and 56.7 μg/kg BW). (A) CHE activity (U/L), (B) direct bilirubin concentration (μmon/L), (C) glucose concentration (mol/L) (* p < 0.05, ** p < 0.01(compared to the control)).

### 3.5. Histopathological examination

No abnormal gross pathology was observed during the dissection. However, compared with the control, treatment-related pathological changes were found only in the liver tissue slices of the ASX exposure groups (Fig. 3). Point steatosis was observed in the 56.7 μg/kg BW ASX exposure group. Necrosis was observed in the 18.9 μg/kg BW and 56.7 μg/kg BW ASX exposure groups.

**Fig.3.**
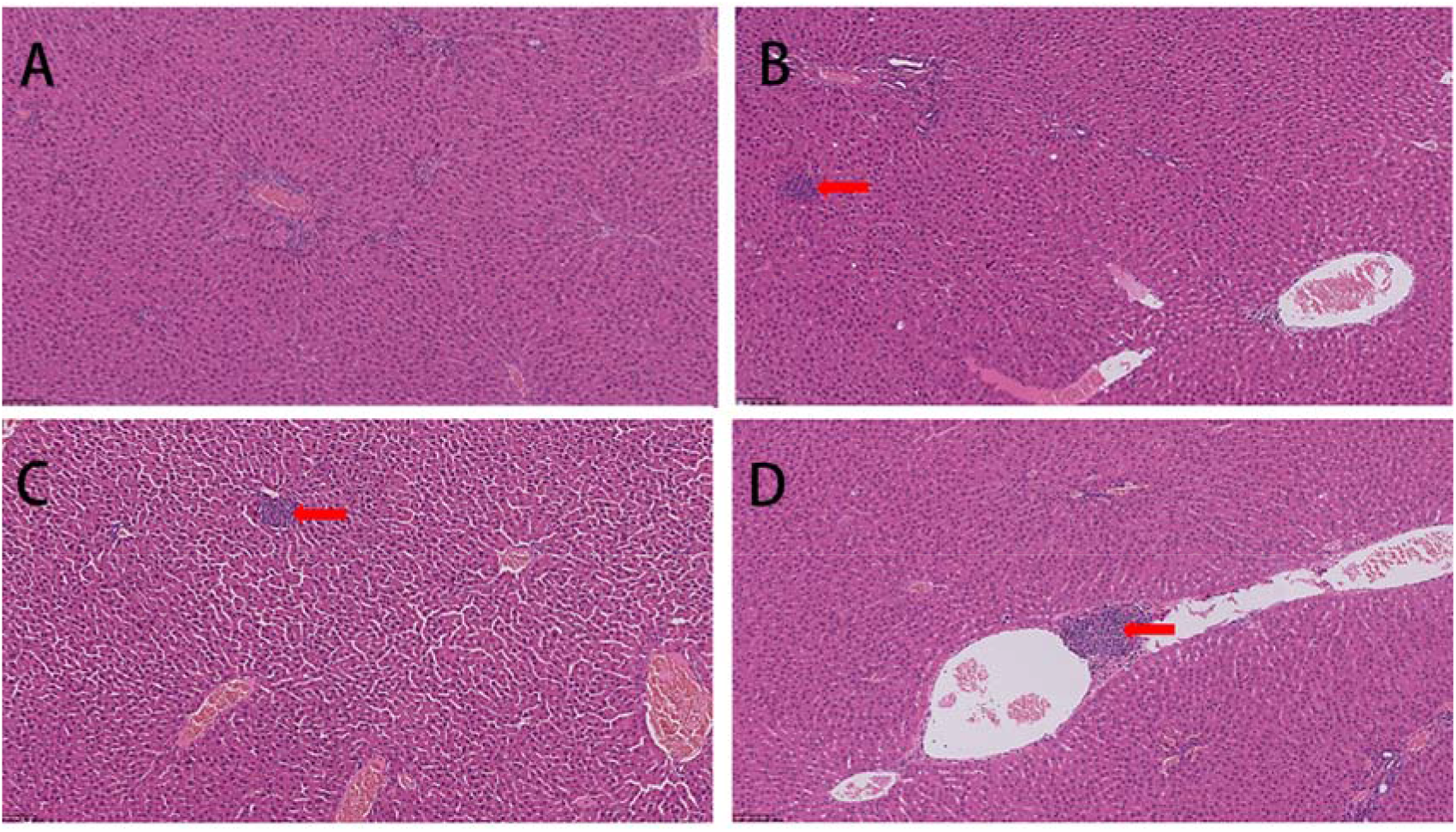
Histopathological HE-stained sections of the rat liver after 12 weeks of oral exposure to ASX (× 200, H&E) A-D: Liver structure of rats; Normal liver in the control group (A); Point necrosis in the 56.7 μg/kg BW group (B); Necrosis and steatosis in the 56.7 μg/kg BW ASX exposure group (C); and Necrosis in the 18.9 μg/kg BW ASX exposure group (D).

## 4. Discussion

The current study aims to calculate the DNEL of ASX in order to protect the general public. The risk to humans can be considered controlled if the estimated exposure levels do not exceed the appropriate DNEL. Furthermore, DNEL can be used to assess exposure and characterize risk. However, there are few data on ASX toxicity. The determination of the ASX safety threshold is critical. The subchronic toxic effects of ASX on rats were investigated using various biochemical, hematological, and histopathological parameters.

In this study, male rats in the middle and high dose groups had higher RBWG by the eighth week, while the exposed rats’ food and water intake did not change significantly. Food and water consumption may be impacted.

Our experiments indicate that the administered doses of ASX (6.3, 18.9, and 56.7 μg/kg) had no statistically significant effect on the observed hematological parameters compared with the control. The observed changes were mostly uncorrelated with the dosages so no reliable conclusions can be drawn from the acquired results.

Histologically, slight pathological alterations in the liver were discovered, which were consistent with the observed biochemical alterations. Detoxification processes take place primarily in the liver, which is the organ that is most exposed to xenobiotics and/or their metabolites. The susceptibility of liver tissues to pesticide-induced stress is determined by the overall balance of oxidative stress and antioxidant capacity (Khan et al., 2005). Carbamates have been linked to liver damage in numerous studies (Brkić et al., 2008; Eraslan et al., 2009; Fujino et al., 2016; Pereira et al., 1991; Wang et al., 2007). CHE activity in female rats was reduced by 9.12%, 9.94%, and 11.64% in the three treated groups, respectively (compared with the control). Cholinesterase inhibition is well known in carbamates (Dési et al., 1974; Padilla et al., 2007; Vandekar et al., 1971; Yu et al., 1972). Serum CHE activity, is an early and sensitive clinical index discovered in patients with anticholinesterase pesticide intoxication (Wang et al., 2014). Cholinesterase activity (pseudocholinesterase) in plasma is a biological indicator of carbamate pesticide exposure (Krechniak and Foss, 1982; Machemer and Pickel, 1994), CHE inhibition is reversible (Gupta et al., 2017), and the degree of acetylcholinesterase inhibition observed at this dosage (24-28%) does not generally cause intoxication symptoms in humans (Wills, 1971/1972). As a result, the CHE inhibition in female rats is insignificant. The total cholesterol level in the three treated groups was significantly lower than in the control group. There have been reports of carbamates lowering total cholesterol levels (Dias et al., 2013). However, some studies have found an increase in serum total cholesterol following carbamate pesticide exposure (Muthuviveganandavel et al., 2008; Oakberg, 1956; Selmanoglu et al., 2001). A significant increase in uric acid was observed in female rats treated with high dose ASX; previous research has shown that carbamate can cause an increase in serum uric acid levels (Nwani et al., 2016). Uric acid is the end product of tissue nucleic acid catabolism, which includes purine and pyrimidine base metabolism. This could be due to purine and pyrimidine degradation or to an increase in uric acid levels caused by either overproduction or the inability to excrete (Wolf, 1972). This suggests that ASX administration may have a negative impact on the kidney. The liver-related parameters CHE activity decreased by 23.72%, 20.31%, and 28.12% in male rats, respectively. Furthermore, a significant increase in direct bilirubin levels was observed in the three treated groups when compared to the control; the same result was observed in a study of rats exposed to Carbofuran (Mashall and Dorough, 1977). Previous research has shown that carbamates such as Carbaryl, Fenoxycarb, Propamocarb, Propoxur, and Carbendazim can cause blood glucose level increases in male rats when given a high dose of ASX (McDaniel et al., 2007; Schmuck and Mihail, 2004; Selmanoglu et al., 2001). A rise in glucose levels could be attributed to a breakdown in glucose intake and utilization by cells.

We, therefore, determined the BMD value based on the findings. To get around the limitations of the NOAEL, which depends on exposure concentration, the EPA suggests that the BMD be estimated. The lowest number was chosen as the BMD value after the BMD computation produced the BMDL, the dosage with a 95% lower confidence level that corresponds to a 10% reaction incidence. Cholesterol level, glucose, uric acid, and CHE activity were the toxicology data for dosage response.

We selected to calculate the BMDL using uric acid, glucose, and cholesterol levels in consideration of the long-term harmful effects, using the exponential and hill models. The BMDL results were respectively 1.07, 2.27, and 0.03 μg/kg BW. As a result, we selected 0.03 μg/kg BW as the Point of Departure (PoD) of DNEL derivation, the dose associated with cholesterol level, and the DNEL was calculated as 0.000375 μg/kg BW by applying an interspecies factor of 4, an intraspecies factor of 10, and an exposure duration factor of 2 as default assessment factors (ECHA, 2012).

## 5. Conclusions

In conclusion, repeated oral ASX exposure for 12 weeks could cause dose-dependent toxic effects in SD rats. Significant changes in blood biochemistry parameters and liver histopathology in response to ASX dose suggested that the liver may be the target organ affected by ASX. Based on the findings, the NOAEL was determined to be less than 6.3 μg/kg BW. As a result, the BMD was calculated to be 0.03 μg/kg BW using dose-response data and a mathematical model. According to information requirements and chemical safety assessment guidelines, the DNEL value converted from these BMD values to human exposure was defined as 0.000375 μg/kg BW.

## Abbreviation

ALB: Albumin
ASX: Aldicarb sulfoxide
ASN: Aldicarb sulfone
ALP: Alkaline phosphate
AST: Aspartate aminotransferase
BASO: Basophil absolute value
BMD: Benchmark dose
BMDL: Benchmark dose low
BWG: Body weight gain
CHE: Cholinesterase
CREA: Creatine
CK: Creatine kinase
DNEL: Derived no-effect level
DBIL: Direct bilirubin
RIVM: Dutch National Institute for Public Health and Environmental
EPA: Environmental protection agency
EOS: Eosinophil absolute value
GLB: Globulin
GGT: Gamm-glutamyl transpeptidase
GLU: Glucose
HCT: Hematocrit
HGB: Hemoglobin
HDLC: High density lipoprotein cholesterol
IBIL: Indirect bilirubin
LDH: Lactate dehydrogenase
P-LCR: Large platelet ratio
LDLC: Low-density lipoprotein cholesterol
LY: Lymphocytes
MCH: Mean hemoglobin account
MCHC: Mean hemoglobin concentration
MPV: Mean platelet volume
MCV: Mean red blood cell volume
MONO: HGB; Monocytes absolute value
NEUT: Neutrophils absolute value
NOAEL: No observed adverse effect level
OECD: Organization for economic cooperation and development
BASO%: Percent basophil
EOS%: Percent eosinophil
LY%: Percent lymphocytes
MONO%: Percent monocytes
NEUT%: Percent neutrophils
PLT: Platelet
PDW: Platelet distribution width
PCV: Platelet packed volume
RBWG: Registration, and authorization of chemicals
REACH: Relative body weight gain;
RBC: Red blood cell
Ratio of white balls: A/G;
RDW-SD: Red blood cell volume distribution width standard deviation
ROW: Relative organ weight
ALT: Serum alanine aminotransferase
Ca: Serum calcium
Cl: Serum chloride
Fe: Serum iron
Mg: Serum magnesium
P: Serum phosphorus
K: Serum potassium
Na: Serum sodium
SD: Sprague-Dawley
TBIL: Total bilirubin
CHOL: Total cholesterol
TP: Total protein
TG: Triglyceride
UA: Uric acid
WBC: White blood cell

## Credit authorship contribution statement

**Yongchao Ji:** Study design, data collection, animal exposure, manuscript draft, data analysis, and interpretation. **Yi Liu:** Data collection and manuscript draft. **Juanjuan Duan:** Data collection. **Yiting Wang:** Data collection. **Yu Wang:** Data collection. **Fan Wang:** Data collection. **Chao Chen:** Manuscript draft. **Wensheng Zhang:** Study design, manuscript draft, data analysis, and interpretation. All the authors listed above have read and approved the version of the final manuscript.

