## supplemental table for "Subchronic oral toxicity study of Aldicarb sulfoxide in Sprague-Dawley rats"

Wensheng Zhang, Ph.D.

Beijing Key Laboratory of Traditional Chinese Medicine Protection and Utilization

Beijing Normal University

No.19, Xinjiekouwai St, Haidian District

Beijing, 100875, China

**Table S1.** Body weight gain in male rats orally exposed to ASX for 12 weeks.

| Concentration (μg/kg BW) | Parameters | week1 | week2 | week3 | week4 | week5 | week6 | week7 | week8 | week9 | week10 | week11 | week12 |
| --- | --- | --- | --- | --- | --- | --- | --- | --- | --- | --- | --- | --- | --- |
| Control (n=10) | RBWG | 0.22±0.06 | 0.39±0.1 | 0.53±0.14 | 0.63±0.15 | 0.74±0.18 | 0.85±0.2 | 0.93±0.22 | 1.0±0.24 | 1.08±0.27 | 1.09±0.33 | 1.12±0.25 | 1.16±0.26 |
| 6.3 (n=10) | RBWG | 0.22±0.02 | 0.38±0.07 | 0.55±0.11 | 0.67±0.15 | 0.77±0.17 | 0.86±0.18 | 0.95±0.2 | 1.02±0.21 | 1.08±0.23 | 1.09±0.22 | 1.06±0.23 | 1.11±0.23 |
|  | % to control | 0% | -3% | 4% | 6% | 4% | 1% | 2% | 2% | 0% | 0% | -5% | -4% |
| 18.9 (n=10) | RBWG | 0.25±0.05 | 0.43±0.1 | 0.59±0.14 | 0.72±0.16 | 0.83±0.18 | 0.95±0.22 | 1.05±0.24 | 1.15±0.26 | 1.23±0.28 | 1.25±0.26 | 1.23±0.27 | 1.27±0.26 |
|  | % to control | 14% | 10% | 11% | 14% | 12% | 12% | 13% | 15% | 14% | 15% | 10% | 9% |
| 56.7 (n=8) | RBWG | 0.24±0.04 | 0.4±0.09 | 0.56±0.15 | 0.68±0.17 | 0.8±0.19 | 0.93±0.21 | 1.01±0.23 | 1.12±0.27 | 1.2±0.27 | 1.24±0.28 | 1.24±0.27 | 1.28±0.27 |
|  | % to control | 9% | 3% | 6% | 8% | 8% | 9% | 9% | 12% | 11% | 14% | 11% | 10% |

* p < 0.05, ** p < 0.01, *** p < 0.001 (compared to the control).

**Table S2.** Body weight gain in female rats orally exposed to ASX for 12 weeks.

| Concentration (μg/kg BW) | Parameters | week1 | week2 | week3 | week4 | week5 | week6 | week7 | week8 | week9 | week10 | week11 | week12 |
| --- | --- | --- | --- | --- | --- | --- | --- | --- | --- | --- | --- | --- | --- |
| Control (n=10) | RBWG | 0.13±0.03 | 0.2±0.03 | 0.31±0.05 | 0.38±0.06 | 0.45±0.07 | 0.49±0.07 | 0.51±0.08 | 0.57±0.07 | 0.6±0.07 | 0.65±0.15 | 0.61±0.08 | 0.66±0.07 |
| 6.3 (n=10) | RBWG | 0.11±0.04 | 0.29±0.19 | 0.36±0.08 | 0.41±0.09 | 0.45±0.12 | 0.5±0.11 | 0.54±0.1 | 0.58±0.12 | 0.62±0.14 | 0.64±0.15 | 0.68±0.15 | 0.68±0.14 |
|  | % to control | -15% | 45% | 16% | 8% | 0% | 2% | 6% | 2% | 3% | -2% | 11% | 3% |
| 18.9 (n=10) | RBWG | 0.12±0.03 | 0.22±0.06 | 0.33±0.07 | 0.4±0.07 | 0.43±0.07 | 0.49±0.08 | 0.54±0.09 | 0.58±0.1 | 0.63±0.11 | 0.63±0.11 | 0.67±0.12 | 0.69±0.14 |
|  | % to control | -8% | 10% | 6% | 5% | -4% | 0% | 6% | 2% | 5% | -3% | 10% | 5% |
| 56.7 (n=10) | RBWG | 0.14±0.03 | 0.18±0.04 | 0.3±0.03 | 0.37±0.05 | 0.42±0.07 | 0.47±0.08 | 0.51±0.1 | 0.56±0.12 | 0.61±0.12 | 0.62±0.15 | 0.65±0.13 | 0.67±0.15 |
|  | % to control | 8% | -10% | -3% | -3% | -7% | -4% | 0% | -2% | 2% | -5% | 7% | 2% |

* p < 0.05, ** p < 0.01, *** p < 0.001 (compared to the control).

**Table S3.** ROW of female rats orally exposed to ASX for 12 weeks

|  | Concentration (μg/kg BW) | | | |
| --- | --- | --- | --- | --- |
| Parameter | Control (n=10) | Low (n=10) | Middle (n=10) | High (n=10) |
| Heart | 0.0036±0.0004 | 0.0035±0.001 | 0.0033±0.0005 | 0.0036±0.0009 |
| Liver | 0.0234±0.0017 | 0.023±0.0032 | 0.0244±0.0052 | 0.0229±0.0036 |
| Spleen | 0.0016±0.0001 | 0.0016±0.0003 | 0.0015±0.0002 | 0.0017±0.0004 |
| Lung | 0.004±0.0005 | 0.004±0.0006 | 0.0039±0.0009 | 0.0043±0.0014 |
| Kidney | 0.0061±0.0005 | 0.0058±0.0013 | 0.0059±0.0008 | 0.0057±0.0025 |
| Adrenal gland | 0.0002±0.0001 | 0.0002±0.0001 | 0.0002±0.0 | 0.0006±0.0012 |
| Uterus ovary | 0.0028±0.0007 | 0.0028±0.0009 | 0.0024±0.0006 | 0.0025±0.0007 |
| Thymus | 0.0015±0.0004 | 0.0014±0.0003 | 0.0013±0.0005 | 0.0015±0.0003 |
| Lymph | 0.002±0.0005 | 0.0023±0.001 | 0.0018±0.0008 | 0.0019±0.0005 |

* p < 0.05, ** p < 0.01, *** p < 0.001 (compared to the control).

**Table S4.** ROW of male rats orally exposed to ASX for 12 weeks

|  | Concentration (μg/kg BW) | | | |
| --- | --- | --- | --- | --- |
| Parameter | Control (n=10) | 6.3 (n=10) | 18.9 (n=10) | 56.7 (n=8) |
| Heart | 0.0033±0.0003 | 0.0036±0.0003 * | 0.0033±0.0004 | 0.0032±0.0003 |
| Liver | 0.022±0.0013 | 0.0178±0.0087 | 0.0228±0.0011 | 0.0228±0.0019 |
| Spleen | 0.0071±0.0181 | 0.0014±0.0002 | 0.0013±0.0002 | 0.0013±0.0001 |
| Lung | 0.0031±0.0004 | 0.0032±0.0004 | 0.0031±0.0004 | 0.0032±0.0004 |
| Kidney | 0.0058±0.0006 | 0.0059±0.0006 | 0.0055±0.0005 | 0.0084±0.0076 |
| Adrenal gland | 0.0001±0.0 | 0.0001±0.0 | 0.0001±0.0 | 0.0001±0.0 |
| Testis | 0.0061±0.0009 | 0.0066±0.0009 | 0.0058±0.0006 | 0.006±0.0005 |
| Epididymis | 0.0024±0.0004 | 0.0026±0.0004 | 0.0023±0.0003 | 0.0026±0.0004 |
| Thymus | 0.001±0.0003 | 0.0009±0.0002 | 0.001±0.0002 | 0.001±0.0002 |
| Lymph | 0.0014±0.0003 | 0.0013±0.0006 | 0.0015±0.0006 | 0.0017±0.0008 |

* p < 0.05, ** p < 0.01, *** p < 0.001 (compared to the control).

**Table S5.** Hematological parameters for female rats after 12 weeks of oral exposure to ASX

|  | Concentration (μg/kg BW) | | | | |
| --- | --- | --- | --- | --- | --- |
| Parameter | | control (n=10) | 6.3 (n=10) | 18.9 (n=10) | 56.7 (n=10) |
| WBC (×10^9^/L) | | 3.56±1.3 | 3.98±1.45 | 4.27±1.71 | 3.28±0.57 |
| LY (%) | | 76.27±3.51 | 78.42±6.32 | 79.69±7.1 | 77.0±6.8 |
| MONO (%) | | 4.84±1.02 | 5.24±0.95 | 4.42±1.45 | 5.12±1.69 |
| NEUT (%) | | 17.04±3.35 | 14.66±5.21 | 14.19±6.34 | 16.13±5.8 |
| EOS (%) | | 1.76±0.6 | 1.64±0.52 | 1.62±0.58 | 1.68±0.74 |
| BASO (%) | | 0.09±0.09 | 0.04±0.05 | 0.08±0.06 | 0.07±0.06 |
| LY (×10^9^/L) | | 2.74±1.1 | 3.13±1.24 | 3.49±1.51 | 2.56±0.61 |
| MONO (×10^9^/L) | | 0.18±0.09 | 0.21±0.09 | 0.18±0.07 | 0.17±0.05 |
| NEUT (×10^9^/L) | | 0.58±0.15 | 0.58±0.27 | 0.53±0.19 | 0.51±0.14 |
| EOS (×10^9^/L) | | 0.06±0.02 | 0.06±0.02 | 0.06±0.03 | 0.05±0.02 |
| BASO (×10^9^/L) | | 0.0±0.0 | 0.0±0.0 | 0.0±0.0 | 0.0±0.0 |
| RBC (×10^12^/L) | | 8.15±0.47 | 8.06±0.39 | 8.02±0.33 | 8.43±0.28 |
| HB (g/L) | | 161.4±7.64 | 159.9±7.97 | 159.2±6.13 | 165.6±7.67 |
| HCT (%) | | 48.51±2.72 | 47.16±2.43 | 47.49±2.08 | 49.2±2.22 |
| MCV (fL) | | 59.5±1.57 | 58.5±1.54 | 59.21±1.68 | 58.4±1.06 |
| MCH (pg) | | 19.82±0.52 | 19.85±0.59 | 19.87±0.61 | 19.62±0.42 |
| MCHC (g/L) | | 332.9±5.34 | 339.2±4.17 * | 335.3±4.05 | 336.2±5.93 |
| RDW-CV (%) | | 13.68±0.52 | 14.3±0.57 * | 13.74±0.34 | 13.92±0.44 |
| RDW-SD (fL) | | 28.46±0.77 | 29.26±1.59 | 28.53±1.05 | 28.37±1.19 |
| PLT (×10^9^/L) | | 1038.3±135.1 | 960.5±122.59 | 1085.9±109.97 | 1078.5±140.09 |
| PDW (fL) | | 15.02±0.24 | 14.88±0.11 | 15.01±0.13 | 14.98±0.18 |
| MPV (fL) | | 7.47±0.71 | 7.1±0.41 | 7.34±0.43 | 7.25±0.56 |
| PCT (%) | | 0.77±0.08 | 0.68±0.11 | 0.8±0.11 | 0.78±0.13 |
| P-LCR (%) | | 11.02±4.78 | 8.14±2.05 | 9.68±2.29 | 9.47±3.19 |

* p < 0.05, ** p < 0.01, *** p < 0.001 (compared to the control).

**Table S6.** Hematological parameters for male rata after 12 weeks of oral exposure to ASX

|  | Concentration (μg/kg BW) | | | | |
| --- | --- | --- | --- | --- | --- |
| Parameter | | control (n=10) | 6.3 (n=10) | 18.9 (n=10) | 56.7 (n=8) |
| WBC (×10^9^/L) | | 6.33±1.16 | 8.53±2.63 * | 6.75±1.84 | 6.5±2.19 |
| LY (%) | | 76.18±3.41 | 72.3±9.36 | 77.83±6.66 | 73.68±9.27 |
| MONO (%) | | 4.54±1.05 | 4.3±2.14 | 4.31±1.28 | 4.62±0.89 |
| NEUT (%) | | 17.35±2.9 | 21.85±8.96 | 16.4±5.65 | 20.04±8.91 |
| EOS (%) | | 1.9±1.13 | 1.52±0.7 | 1.41±0.5 | 1.64±0.65 |
| BASO (%) | | 0.03±0.05 | 0.03±0.05 | 0.05±0.05 | 0.02±0.04 |
| LY (×10^9^/L) | | 4.82±0.88 | 6.24±2.29 | 5.29±1.61 | 4.89±1.95 |
| MONO (×10^9^/L) | | 0.28±0.07 | 0.34±0.13 | 0.3±0.14 | 0.31±0.14 |
| NEUT (×10^9^/L) | | 1.11±0.31 | 1.82±0.88 * | 1.07±0.44 | 1.2±0.43 |
| EOS (×10^9^/L) | | 0.12±0.07 | 0.12±0.06 | 0.1±0.05 | 0.1±0.03 |
| BASO (×10^9^/L) | | 0.0±0.0 | 0.0±0.0 | 0.0±0.0 | 0.0±0.0 |
| RBC (×10^12^/L) | | 8.78±0.53 | 8.88±0.35 | 8.57±0.48 | 8.54±0.15 |
| HB (g/L) | | 163.9±7.56 | 167.6±5.54 | 161.5±6.14 | 159.75±5.36 |
| HCT (%) | | 49.37±1.97 | 49.22±1.69 | 48.34±1.73 | 48.42±2.01 |
| MCV (fL) | | 56.33±1.62 | 55.49±2.06 | 56.53±2.74 | 56.68±1.9 |
| MCH (pg) | | 18.7±0.43 | 18.9±0.73 | 18.87±0.88 | 18.71±0.43 |
| MCHC (g/L) | | 332.2±7.22 | 340.4±3.83 * | 334.0±5.27 | 330.38±5.52 |
| RDW-CV (%) | | 14.93±0.38 | 14.93±0.44 | 15.0±0.58 | 14.94±0.53 |
| RDW-SD( fL) | | 29.28±0.51 | 28.83±0.88 | 29.53±0.68 | 29.52±0.47 |
| PLT (×10^9^/L) | | 1071.3±108.67 | 1097.0±111.39 | 1052.2±267.9 | 1033.12±85.6 |
| PDW (fL) | | 14.92±0.17 | 14.87±0.1 | 14.9±0.22 | 14.85±0.11 |
| MPV (fL) | | 7.1±0.37 | 7.21±0.41 | 7.06±0.4 | 7.08±0.38 |
| PCT (%) | | 0.76±0.11 | 0.79±0.1 | 0.74±0.19 | 0.73±0.07 |
| P-LCR (%) | | 8.9±1.88 | 9.33±2.17 | 8.73±2.33 | 8.62±1.93 |

* p < 0.05, ** p < 0.01, *** p < 0.001 (compared to the control).

**Table S7.** Biochemical parameters for female rats after 12 weeks of oral exposure to ASX.

|  | Concentration (μg/kg BW) | | | | |
| --- | --- | --- | --- | --- | --- |
| Parameter | | control (n=10) | 6.3 (n=10) | 18.9 (n=10) | 56.7 (n=10) |
| AST (U/L) | | 175.13±45.35 | 131.1±43.52 | 158.19±73.52 | 186.76±73.2 |
| ALT (U/L) | | 40.5±4.64 | 30.75±7.61 | 40.58±23.2 | 43.81±16.18 |
| TBIL (μmol/L) | | 2.05±0.34 | 1.88±0.37 | 1.99±0.41 | 2.18±0.56 |
| DBIL (μmol/L) | | 1.38±0.29 | 1.32±0.33 | 1.28±0.18 | 1.37±0.33 |
| IBIL (μmol/L) | | 0.68±0.29 | 0.56±0.31 | 0.71±0.28 | 0.81±0.42 |
| TP (g/L) | | 70.86±4.0 | 63.13±7.54 | 65.64±12.17 | 70.02±4.43 |
| ALB (g/L) | | 30.53±2.15 | 26.49±3.34 | 28.12±4.77 | 29.31±1.75 |
| GLB (g/L) | | 40.33±2.31 | 36.64±4.67 | 37.52±7.51 | 40.71±2.94 |
| A/G | | 0.76±0.04 | 0.73±0.06 | 0.76±0.05 | 0.72±0.03 |
| CHE (U/L) | | 4173.75±130.82 | 3793.25±256.26** | 3758.88±241.31*** | 3688.38±304.87*** |
| GGT (U/L) | | 0.01±0.03 | 0.04±0.07 | 0.08±0.08 | 0.11±0.11* |
| ALP (U/L) | | 38.56±6.32 | 38.24±12.25 | 36.36±9.99 | 42.39±10.16 |
| UREA (μmol/L) | | 13.21±2.41 | 14.19±4.1 | 13.97±3.89 | 14.4±3.25 |
| Creatinine (μmol/L) | | 45.7±6.12 | 51.0±11.66 | 46.2±16.36 | 46.1±10.2 |
| UA (μmol/L) | | 116.25±18.97 | 111.62±37.93 | 94.0±24.28 | 138.75±19.98* |
| GLU (mol/L) | | 8.27±1.88 | 8.13±2.37 | 7.24±2.1 | 7.46±2.02 |
| TG (mol/L) | | 0.23±0.1 | 0.11±0.03 | 0.21±0.1 | 0.34±0.18 |
| CHOL (mol/L) | | 2.23±0.26 | 1.50±0.14** | 1.88±0.2* | 1.79±0.21** |
| HDLC (mol/L) | | 0.53±0.04 | 0.43±0.06 | 0.48±0.1 | 0.5±0.05 |
| LDLC (mol/L) | | 0.27±0.05 | 0.19±0.07* | 0.25±0.09 | 0.22±0.04* |
| CK( U/L) | | 2132.64±940.57 | 1355.68±527.04 | 1249.54±488.19 | 2096.62±486.85 |
| LDH (U/L) | | 1616.8±567.21 | 1421.2±633.83 | 1526.9±666.46 | 1840.8±564.22 |
| K (mmol/L) | | 5.52±0.6 | 5.0±0.55 | 5.06±0.82 | 5.48±0.33 |
| Na (mmol/L) | | 146.99±1.65 | 142.43±11.01 | 142.58±18.46 | 145.85±1.44 |
| Cl (mmol/L) | | 106.96±2.27 | 105.1±7.68 | 103.26±11.71 | 106.27±1.79 |
| Ca (mmol/L) | | 2.22±0.1 | 2.14±0.27 | 2.11±0.4 | 2.2±0.19 |
| Mg (mmol/L) | | 1.36±0.19 | 1.27±0.23 | 1.23±0.27 | 1.37±0.08 |
| P (mmol/L) | | 3.8±0.98 | 4.05±0.73 | 3.8±0.98 | 3.81±0.67 |
| Fe (mol/L) | | 37.43±3.96 | 37.61±10.15 | 36.87±7.58 | 37.8±5.0 |

* p < 0.05, ** p < 0.01, *** p < 0.001 (compared to the control).

**Table S8.** Biochemical parameters for male rats after 12 weeks of oral exposure to ASX.

|  | Concentration (μg/kg BW) | | | | |
| --- | --- | --- | --- | --- | --- |
| Parameter | | Control (n=10) | 6.3 (n=10) | 18.9 (n=10) | 56.7 (n=8) |
| AST (U/L) | | 157.62±48.9 | 133.95±27.14 | 164.64±41.78 | 182.3±133.8 |
| ALT (U/L) | | 47.15±25.63 | 33.56±7.02 | 40.66±8.09 | 63.61±66.11 |
| TBIL (μmol/L) | | 1.85±0.31 | 2.04±0.51 | 2.19±0.61 | 2.0±0.39 |
| DBIL (μmol/L) | | 1.29±0.18 | 1.71±0.27** | 1.61±0.27** | 1.72±0.46** |
| IBIL (μmol/L) | | 0.52±0.24 | 0.41±0.32 | 0.56±0.47 | 0.31±0.23 |
| TP (g/L) | | 54.3±10.55 | 53.56±9.25 | 59.63±1.31 | 57.34±2.83 |
| ALB (g/L) | | 20.31±3.93 | 20.25±3.3 | 21.91±0.53 | 21.42±1.41 |
| GLB (g/L) | | 33.99±6.74 | 33.31±6.11 | 37.72±1.23 | 35.91±2.38 |
| A/G | | 0.6±0.03 | 0.61±0.05 | 0.58±0.02 | 0.6±0.05 |
| CHE (U/L) | | 2664.88±478.83 | 2032.73±459.75* | 2204.92±311.98* | 1915.43±622.73* |
| GGT (U/L) | | 0.11±0.25 | 0.01±0.03 | 0.28±0.71 | 0.02±0.07 |
| ALP (U/L) | | 85.08±24.06 | 81.51±20.58 | 91.17±13.06 | 85.34±13.68 |
| UREA (μmol/L) | | 10.62±1.95 | 11.7±1.46 | 12.09±1.81 | 12.21±2.05 |
| Creatinine (μmol/L) | | 40.9±14.05 | 32.3±6.75 | 33.8±7.17 | 40.38±6.16 |
| UA (μmol/L) | | 128.9±39.26 | 117.8±32.09 | 112.4±39.53 | 140.62±29.6 |
| GLU (mol/L) | | 9.53±2.07 | 8.36±1.15 | 10.74±2.83 | 13.85±3.33* |
| TG (mol/L) | | 0.22±0.11 | 0.29±0.13 | 0.41±0.26 | 0.39±0.31 |
| CHOL (mol/L) | | 1.44±0.34 | 1.49±0.3 | 1.75±0.25 | 1.61±0.29 |
| HDLC (mol/L) | | 0.3±0.08 | 0.28±0.04 | 0.34±0.07 | 0.32±0.05 |
| LDLC (mol/L) | | 0.29±0.09 | 0.31±0.07 | 0.34±0.07 | 0.32±0.08 |
| CK (U/L) | | 2196.99±749.18 | 1731.51±606.44 | 1977.53±1313.04 | 1854.98±991.68 |
| LDH (U/L) | | 1732.2±522.69 | 1649.7±432.54 | 1831.6±456.53 | 1774.5±853.24 |
| K (mmol/L) | | 5.54±0.76 | 5.35±0.59 | 5.63±0.37 | 5.48±0.59 |
| Na (mmol/L) | | 140.26±17.35 | 138.3±14.95 | 148.09±1.64 | 146.52±3.25 |
| Cl (mmol/L) | | 104.14±11.35 | 101.88±10.36 | 107.82±2.03 | 105.81±3.71 |
| Ca (mmol/L) | | 1.92±0.38 | 1.78±0.26 | 2.03±0.09 | 2.05±0.37 |
| Mg (mmol/L) | | 1.09±0.23 | 1.08±0.19 | 1.16±0.08 | 1.17±0.12 |
| P (mmol/L) | | 3.53±0.64 | 3.43±0.61 | 3.66±0.37 | 3.75±0.59 |
| Fe (mol/L) | | 17.93±6.48 | 17.39±4.8 | 18.4±3.52 | 17.34±4.94 |

* p < 0.05, ** p < 0.01, *** p < 0.001 (compared to the control).
